# Unpredictable effects of the genetic background of transgenic lines in physiological quantitative traits

**DOI:** 10.1101/494419

**Authors:** Amalia Evangelou, Anastasia Ignatiou, Chloi Antoniou, Sofia Kalanidou, Sotiroula Chatzimatthaiou, Gavriella Shianiou, Soteroulla Ellina, Rafaella Athanasiou, Myrofora Panagi, Yiorgos Apidianakis, Chrysoula Pitsouli

## Abstract

Physiological, fitness and disease phenotypes are complex traits exhibiting continuous variation in natural populations. To understand complex trait gene functions transgenic lines of undefined genetic background are often combined to assess quantitative phenotypes ignoring the impact of genetic polymorphisms. Here, we used inbred wild-type strains of the *Drosophila* Genetics Reference Panel to assess the phenotypic variation of six physiological and fitness traits, namely, female fecundity, survival and intestinal mitosis upon oral infection, defecation rate and fecal pH upon oral infection, and terminal tracheal cell branching in hypoxia. We found continuous variation in the approximately 150 strains tested for each trait, with extreme values differing by more than four standard deviations for all traits. In addition, we assessed the effects of commonly used *Drosophila UAS-RNAi* transgenic strains and their backcrossed isogenized counterparts, in the same traits plus baseline intestinal mitosis and tracheal branching in normoxia, in heterozygous conditions, when only half of the genetic background was different among strains. We tested 20 non-isogenic strains (10 KK and 10 GD) from the Vienna *Drosophila* Resource Center and their isogenized counterparts without Gal4 induction. Survival upon infection and female fecundity exhibited differences in 50% and 40% of the tested isogenic vs. non-isogenic pairs, respectively, whereas all other traits were affected in only 10-25% of the cases. Upon Gal4-induced *UAS-RNAi* expression, 4 out of 11 isogenic vs. non-isogenic pairs tested exhibited differences in survival to infection. And *vice versa*, crossing a single *UAS-RNAi* line with a Gal4 transgene inserted in different genetic backgrounds exhibited quantitative variations that were unpredictable on the basis of pure line performance. Thus, irrespective of the trait of interest, the genetic background of commonly used transgenic strains needs to be considered carefully during experimentation.

## INTRODUCTION

Physiological, morphological, biochemical, behavioral, fitness and disease phenotypes exhibit continuous variation among natural populations and are therefore considered complex or quantitative traits. This continuity in variation is attributed to a combination of genetic composition, environmental effects and gene-environment interactions (Mackay, 2009). To assess the impact of environmental factors on defined genetic backgrounds to tease out the effect of the genotype from that of the environment, inbred lines can be used. Naturally-occurring phenotypic variation in distinct inbred backgrounds allows the implementation of genome-wide association (GWA) and the identification of key genetic variants impinging on the phenotype under study. The *Drosophila* Genetics Reference Panel (DGRP) collection corresponds to a set of >200 inbred sequenced fruit fly strains, which facilitate GWA studies (Mackay et al., 2012; Huang et al., 2014,). GWAS analyses using the DGRP collection have identified significant genomic associations with a variety of fitness and physiology traits ranging from lifespan and sleep to abdominal pigmentation (Durham et al., 2014; Ivanov et al., 2015; Harbison et al., 2013; Dembeck et al., 2015; Sunaga et al., 2016).

The natural variation of the wild-type genetic background not only affects complex traits, but it often also affects the expressivity and penetrance of different mutations. Genetic background effects have been observed in all model organisms tested, including fruit flies, mice, nematodes, yeast, plants and bacteria (Chandler et al., 2013). These effects potentially led to contradictory results across studies that investigated the role of genes affecting longevity, such as *Indy* and *sir-2*, whereby the original papers that implicated the genes in increased longevity (Rogina et al., 2000; Rogina et al., 2004) could not be replicated in subsequent studies performed in different wild-type genetic backgrounds (Toivonen et al., 2007; Burnett et al., 2011; Viswanathan and Guarente, 2011). Furthermore, extensive studies assessing the phenotype of various alleles of the gene *sd* in *Drosophila*, which affect wing size, have shown that the same allele is able to generate a continuum of phenotypes when introduced in different wild-type genetic backgrounds (Chari and Dworkin, 2013; Dworkin et al., 2009). In addition, it has been shown that the genetic background can also affect RNAi phenotypes. For example, injection of RNAi constructs in different wild-type genetic backgrounds of *Tribolium* led to distinct phenotypes, which were attributed not to off-target effects or differences in the function of the RNAi machinery, but to maternal effects (Kitzman et al., 2013). In conclusion, such phenotypic inconsistencies underscore the importance of controlling for the genetic background in similar studies.

Backcrossing is an established breeding scheme where a characteristic (i.e. a trait, a gene, a locus or a chromosome segment) is introgressed from a donor parent into the genomic background of a recurrent parent. Progeny of successive generations are selected for the characteristic of interest and subsequently backcrossed to the recurrent parent. As backcrossed generations accumulate, the proportion of the genome of the donor parent tends to zero, except of the part hosting the characteristic of interest. Backcrossing and introgression have long been used in breeding improvement programs, but backcrossing is also particularly useful in dissecting the genetic architecture of quantitative traits, because it can isolate a gene or chromosomal locus of interest in different genetic backgrounds (Hospital, 2005). Isogenic (or congenic) strains derive from backcrossing schemes and contain nearly identical genomes; namely, the genome of the recurrent parent except from the small part that is selected to differ (Kooke et al., 2012). Isogenic strains incorporating specific transgenes (with P or other transposable element ends or phage ends) in different genetic backgrounds can be easily generated in *Drosophila* using phenotypic markers of transgenesis (i.e. eye color, fluorescence) as a tool for selection.

Here, we used *Drosophila melanogaster* to assess the effects of the genetic background in six different fitness and physiology traits (female fecundity, survival and intestinal stem cell mitosis upon oral bacterial infection, defecation rate and fecal pH upon oral bacterial infection, and tracheal branching upon hypoxia). First, we assessed natural phenotypic variation of these traits using the DGRP collection. Then, we modified the genetic background of *UAS-RNAi* transgenic lines and compared the acquired phenotypes in isogenic vs. non-isogenic lines in the absence and presence of a Gal4 driver. Finally, we modified the genetic background of a Gal4 driver and assessed its phenotypic consequences. We found that the genetic background affects different traits to different extents and we propose ways to control for such effects in future experiments.

## RESULTS

### The genetic background plays a key role to the phenotype of complex traits

To understand the complexity of various quantitative fitness and physiology traits of interest, we screened the DGRP collection of wild-type inbred isogenic strains (Mackay et al., 2012) for female fecundity, survival upon oral bacterial infection, intestinal physiology upon oral bacterial infection (intestinal stem cell mitosis-mediated regeneration, defecation rate and fecal pH) and cellular response to hypoxia (terminal tracheal branching). All traits were assessed in *Drosophila* adults, except tracheal branching, which was assessed in third instar larvae. Z-score analysis underscored the continuity of the observed phenotypes for each trait (Figure 1). The standard deviation range of the screened DGRP strains, which indicates the phenotypic range, for each screen equaled 5.57, 5.24, 5.43, 4.28, 4.55 and 4.85 for fecundity, survival, intestinal mitosis, defecation rate, fecal pH and branching, respectively. GWA analysis of each screen via the DGRP2 algorithm (http://dgrp2.gnets.ncsu.edu/) identified 36, 44, 95, 32, 18 and 21 variants associated with fecundity, survival, intestinal mitosis, defecation rate, fecal pH and branching, respectively. To assess if the six traits tested are functionally linked, we performed 15 pair-wise correlation analyses of each trait values for a common set of 128 DGRP strains. A weak correlation was found between the defecation rate and fecal pH (R^2^=0.1345, *p*=0.1) (Fig. 2L) with all other comparisons exhibiting essentially no correlation (R^2^<0.04) (Fig. 2A-K, M-O). Thus, no significant correlation was observed between traits (Figure 2), and the six different traits can be considered largely independent.

**Figure 1:**
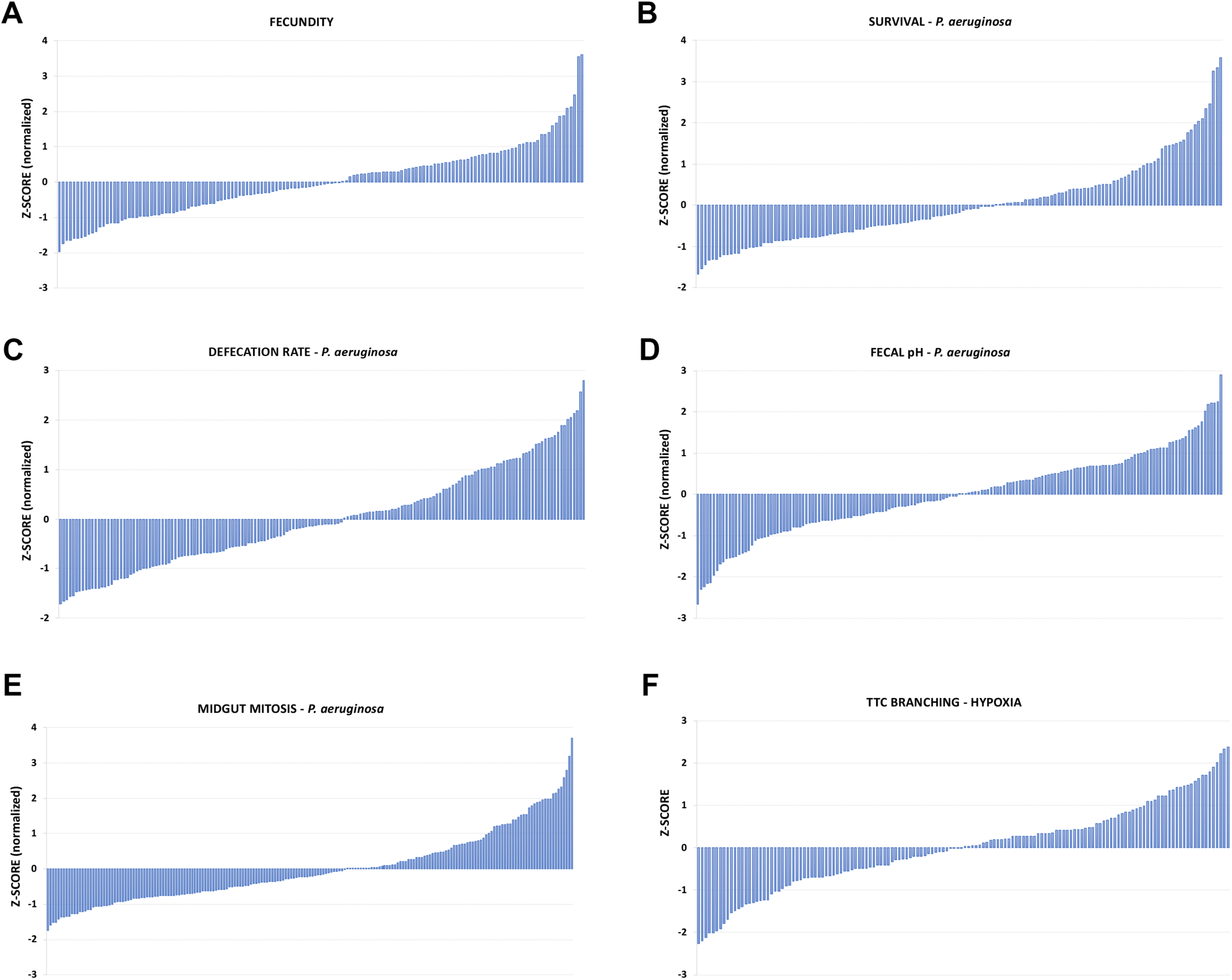
Z-score analysis of the DGRP phenotypes of six complex traits. (**A**) Female fecundity, (**B**) Survival upon oral infection with *P. aeruginosa*, (**C**) Defecation rate upon oral infection with *P. aeruginosa*, (**D**) Fecal pH upon oral infection with *P. aeruginosa*, (**E**) Intestinal mitosis upon oral infection with *P. aeruginosa*, and (**F**) Terminal tracheal branching in hypoxia.

**Figure 2:**
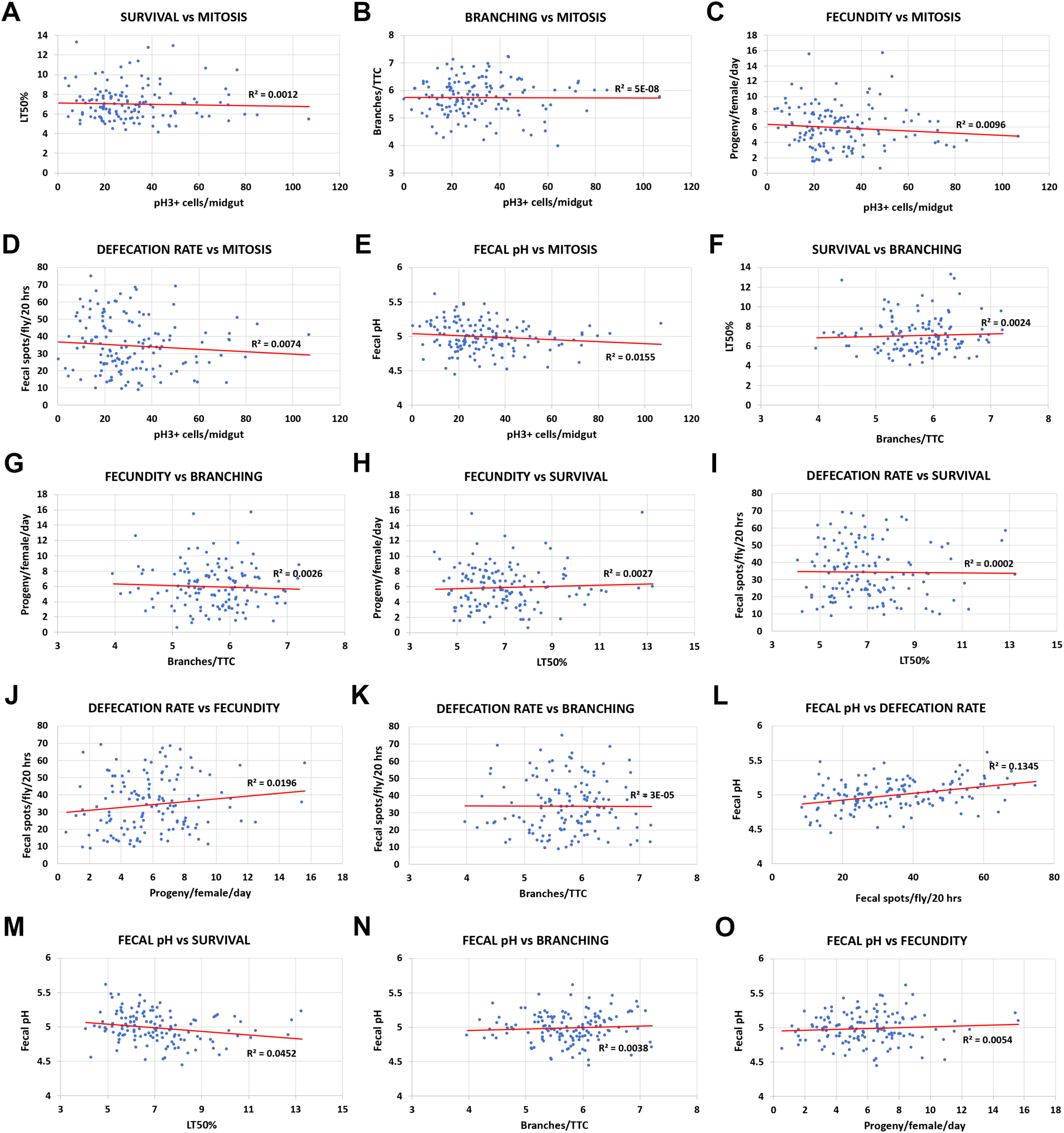
Pair-wise correlation analyses of the six DGRP screens. (**A**) Survival vs intestinal mitosis upon oral infection with *P. aeruginosa*. (**B**) Branching in hypoxia vs intestinal mitosis upon oral infection with *P. aeruginosa*. (**C**) Fecundity vs intestinal mitosis upon oral infection with *P. aeruginosa*. (**D**) Defecation rate upon oral infection with *P. aeruginosa* vs intestinal mitosis upon oral infection with *P. aeruginosa*. (**E**) Fecal pH upon oral infection with *P. aeruginosa* vs intestinal mitosis upon oral infection with *P. aeruginosa*. (**F**) Survival upon oral infection with *P. aeruginosa* vs branching in hypoxia. (**G**) Fecundity vs branching in hypoxia. (**H**) Fecundity vs survival upon oral infection with *P. aeruginosa*. (**I**) Defecation rate upon oral infection with *P. aeruginosa* vs survival upon oral infection with *P. aeruginosa*. (**J**) Defecation rate upon oral infection with *P. aeruginosa* vs fecundity. (**K**) Defecation rate upon oral infection with *P. aeruginosa* vs branching in hypoxia. (**L**) Fecal pH upon oral infection with *P. aeruginosa* vs defecation rate upon oral infection with *P. aeruginosa*. (**M**) Fecal pH upon oral infection with *P. aeruginosa* vs survival upon oral infection with *P. aeruginosa*. (**N**) Fecal pH upon oral infection with *P. aeruginosa* vs branching in hypoxia. (**O**) Fecal pH upon oral infection with *P. aeruginosa* vs fecundity. Trendlines are indicated in red and the R^2^ values are shown in all graphs.

### The genetic background of UAS-RNAi transgenes (without Gal4) affects complex traits to different extents

Since the effect of the homozygous wild-type genotype produces distinct quantitative phenotypes in all traits studied, we devised a way to assess the effects of heterozygosis in the above traits. The use of heterozygotes is necessary in most genetic experiments utilizing the classical genetic tools available in *Drosophila*, such as the Gal4-UAS system (Brand and Perrimon, 1993) and the ever-increasing number of *UAS-RNAi* lines targeting practically any gene of the fly genome. Initially, to avoid the effect of overexpression by the Gal4, we decided to assess the effect of the genetic background on the *UAS-RNAi* insertion alone. All *UAS-RNAi* lines were generated in an isogenic *w^1118^* host strain at the VDRC and the balancer stocks used to map or stabilize insertions were also isogenic to the VDRC *w^1118^* strain. Therefore, and to maintain the insertions as closely as possible to their original genotype, we introgressed the *UAS-RNAi* insertions of twenty randomly-selected VDRC lines in our laboratory *w^1118^* strain for further experiments. Ten strains from the VDRC KK library, generated via site-specific *φχ31* recombination (Markstein et al., 2008), and ten strains from the VDRC GD library (Dietzl et al., 2007), generated via P-element random transposition (Spradling and Rubin, 1982), targeting in pairs the same gene were used. To generate “isogenic” *UAS-RNAi* strains, we backcrossed each VDRC KK and GD strain (from now on called “non-isogenic”) to our laboratory *w^1118^* strain for at least six generations. During backcrossing the fraction of the donor parent (VDRC line) halves after every backcross: 50%, 25%, 12.5% etc., in F1, 1^st^ backcross, 2^nd^ backcross etc., respectively. The average proportion of the recurrent parental genome after each backcross increases and can be calculated by the formula: (2^(b+1)^ − 1)/2^(b+1)^ or 1 − (1/2)^(b+1)^, where b equals the number of backcrosses assuming an infinite population. Thus, the proportion of the donor genome is given as (1/2)^(b+1)^ (Kooke et al., 2012). Assuming six generations of backcrossing, the recurrent parental genome (our laboratory *w^1118^*) equals 99.21% and the donor genome (VDRC *w^1118^*) 0.79%.

The homozygous non-isogenic (VDRC) and isogenic (VDRC in our *w^1118^* strain) lines were subsequently crossed to *w^1118^*, the heterozygous progeny of the crosses were collected and assessed for eight fitness and physiology traits. In addition to the traits studied in the DGRP screening, we also tested intestinal mitosis during homeostasis (without infection) and tracheal branching in normoxia (21% O_2_). GD and KK non-isogenic and isogenic lines were assessed in pairs and those producing significantly different phenotypes were quantified (Figure 3). In this scheme, the progeny of the non-isogenic cross to our laboratory *w^1118^* contain 50% of our laboratory *w^1118^* genome and 50% of the VDRC *w^1118^* genome, whereas the progeny of the isogenic cross to our laboratory *w^1118^* contain approximately 100% of our laboratory *w^1118^* genome. Therefore, the genome of the progeny of the two crosses differs by up to 50%. Theoretically, since the VDRC lines were generated in a *w^1118^* genetic background, we expected not to find significant differences between the original VDRC strains (non-isogenic) and the introgressed lines (isogenic). Strikingly, we observed different impact of isogenization in the various traits and this was independent of the type of transgene (GD, inserted in a random genomic location; or KK, inserted in a specific genomic location); the most affected being survival and fecundity, with 50% and 40% of the isogenic vs. non-isogenic pairs producing significantly different phenotypes (Figure 3A-D), respectively, and the least affected being tracheal branching in normoxia and hypoxia with only 10% of the isogenic vs. non-isogenic pairs producing significantly different phenotypes (Figure 3M-P).

**Figure 3:**
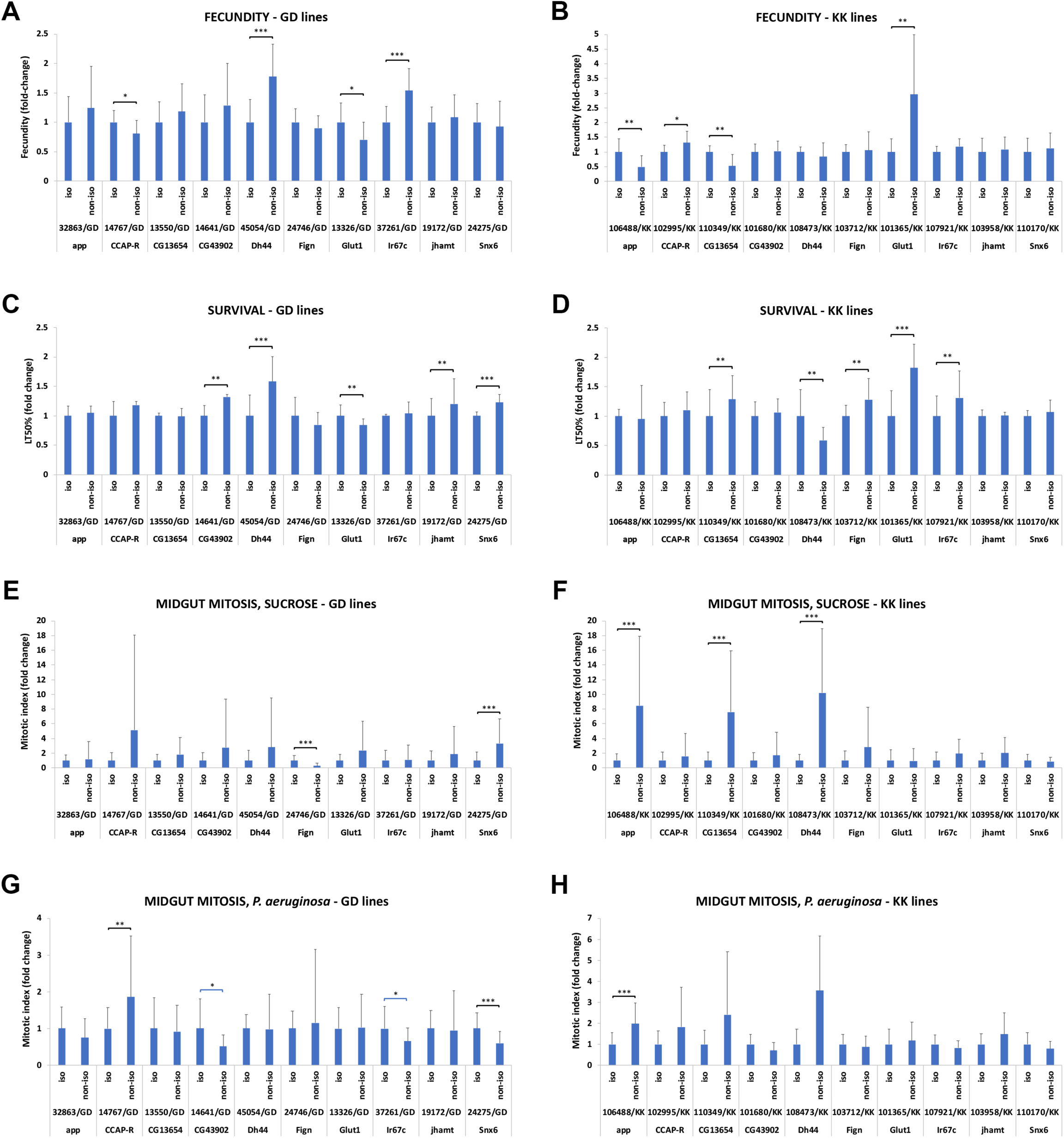
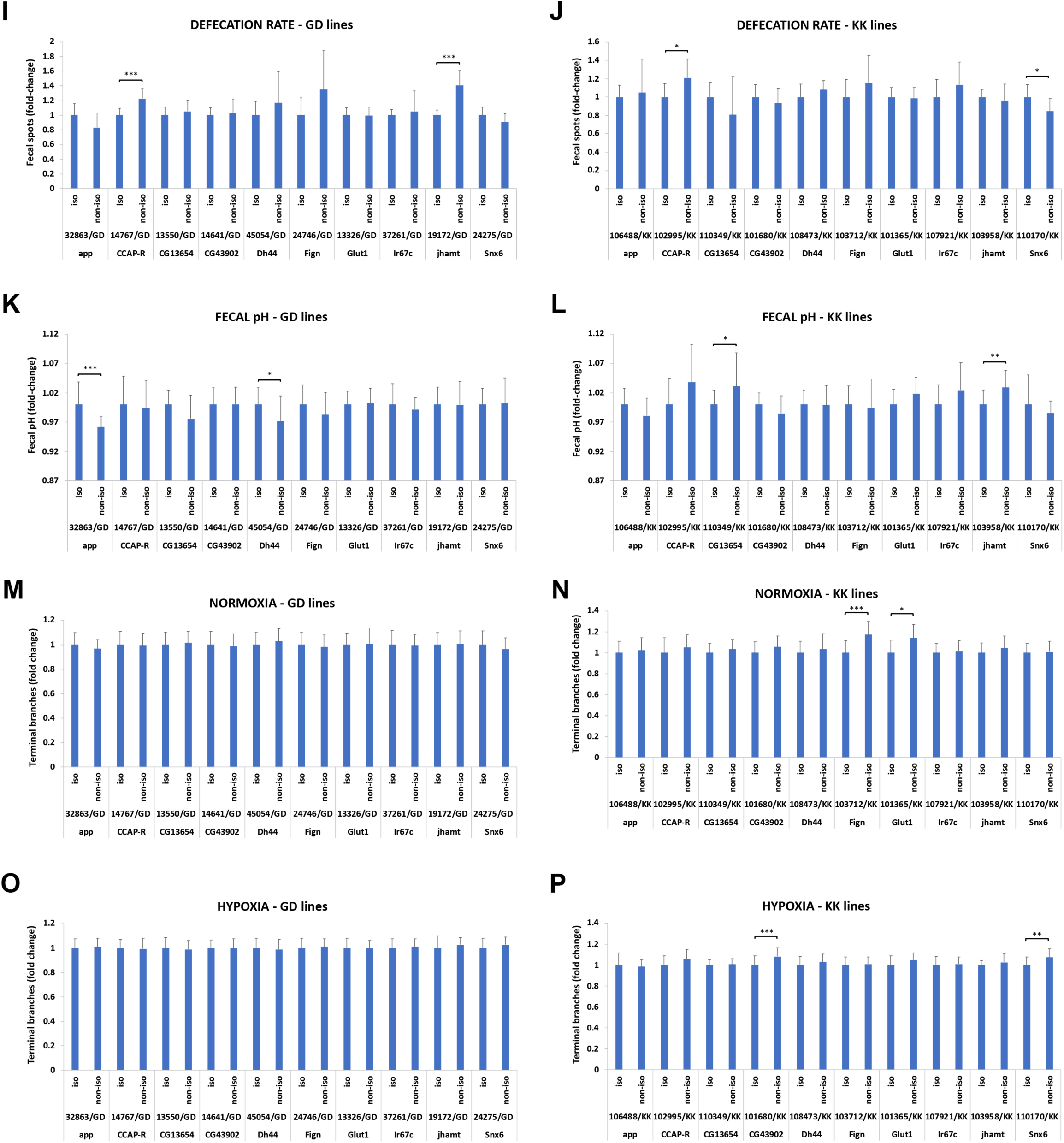
Isogenic vs non-isogenic comparisons of twenty *UAS-RNAi* transgenic lines in the absence of Gal4 for eight traits. (**A-B**) Fecundity in GD and KK lines. (**C-D**) Survival upon oral infection with *P. aeruginosa* in GD and KK lines. (**E-F**) Midgut mitosis in uninfected conditions in GD and KK lines. (**G-H**) Midgut mitosis upon oral infection with *P. aeruginosa* in GD and KK lines. (**I-J**) Defecation rate upon oral infection with *P. aeruginosa* in GD and KK lines. (**K-L**) Fecal pH upon oral infection with *P. aeruginosa* in GD and KK lines. (**M-N**) Tracheal branching in normoxia in GD and KK lines. (**O-P**) Tracheal branching in hypoxia in GD and KK lines. Significant pair-wise comparisons are indicated with brackets, whereby * 0.01<*p*≤0.05; ** 0.001<*p*≤0.01; *** *p*≤0.001.

Specifically, 4 out of 10 GD (2 up, 2 down) and 4 out of 10 KK (2 up, 2 down) isogenic vs. non-isogenic pairs (40% in total) differed in fecundity (Figure 3A-B); 5 out of 10 GD (4 down, 1 up) and 5 out of 10 KK (4 down, 1 up) isogenic vs. non-isogenic pairs (50% in total) differed in survival upon pathogenic oral bacterial infection (Figure 3C-D); 2 out of 10 GD (1 up, 1 down) and 3 out of 10 KK (3 down) isogenic vs. non-isogenic pairs (25% in total) differed in intestinal mitosis in homeostatic conditions (Figure 3E-F); 4 out of 10 GD (1 down, 3 up) and 1 out of 10 KK (1 down) isogenic vs. non-isogenic pairs (25% in total) differed in intestinal mitosis upon oral pathogenic infection (Figure 3G-H); 2 out of 10 GD (2 down) and 2 out of 10 KK (1 up, 1 down) isogenic vs. non-isogenic pairs (20% in total) differed in defecation rate, i.e. number of fecal spots upon bacterial infection (Figure 3I-J); 2 out of 10 GD (2 up) and 2 out of 10 KK (2 down) isogenic vs. non-isogenic pairs (20% in total) differed in fecal pH upon bacterial infection (Figure K-L); 0 out of 10 GD and 2 out of 10 KK (2 down) isogenic vs. non-isogenic pairs (10% in total) differed in tracheal branching in normoxia (Figure 3M-N); and 0 out of 10 GD and 2 out of 10 KK (2 down) isogenic vs. non-isogenic pairs (10% in total) differed in tracheal branching in hypoxia (Figure 3O-P). A summary of these data is presented in Table 1. In conclusion, the genetic background affected each complex trait more or less frequently.

**Table 1:** Isogenic and non-isogenic GD and KK transgenic *UAS-RNAi* lines present variable differences in various phenotypic traits in the absence of Gal4. The last column compares the isogenic and non-isogenic strains with regards to the direction of the phenotype (up: increased, and down: decreased in the isogenic when compared to the non-isogenic).

|  | Isogenic vs non-isogenic pairs with difference (20 total) (% of total) | GD pairs (10 total) | Up in GD isogenic |
| --- | --- | --- | --- |
|  |  |  | Down in GD isogenic |
|  |  | KK pairs (10 total) | Up in KK isogenic |
|  |  |  | Down in KK isogenic |
| Fecundity | 8 (40%) | 4 | 2 |
|  |  |  | 2 |
|  |  | 4 | 2 |
|  |  |  | 2 |
| Survival upon oral infection | 10 (50%) | 5 | 1 |
|  |  |  | 4 |
|  |  | 5 | 1 |
|  |  |  | 4 |
| Midgut mitosis in baseline conditions | 5 (25%) | 2 | 1 |
|  |  |  | 1 |
|  |  | 3 | 0 |
|  |  |  | 3 |
| Midgut mitosis upon oral infection | 5 (25%) | 4 | 1 |
|  |  |  | 3 |
|  |  | 1 | 0 |
|  |  |  | 1 |
| Defecation rate upon oral infection | 4 (20%) | 2 | 0 |
|  |  |  | 2 |
|  |  | 2 | 1 |
|  |  |  | 1 |
| Fecal pH upon oral infection | 4 (20%) | 2 | 2 |
|  |  |  | 0 |
|  |  | 2 | 0 |
|  |  |  | 2 |
| Terminal branching in normoxia | 2 (10%) | 0 | 0 |
|  |  |  | 0 |
|  |  | 2 | 0 |
|  |  |  | 2 |
| Terminal branching in hypoxia | 2 (20%) | 0 | 0 |
|  |  |  | 0 |
|  |  | 2 | 0 |
|  |  |  | 2 |

Further analysis of the differences of isogenic vs. non-isogenic pairs in the various assays indicated that, although statistically significant, the phenotype might differ by a small or large percentage, as reflected by the fold-change difference between the genotypes. For example, for the total of 10 isogenic/non-isogenic pairs exhibiting differences in survival upon infection, the percentage of phenotypic difference ranges from 16% (13326/GD iso/non-iso pair) to 81.8% (101365/KK iso/non-iso pair). Since the biggest difference between an isogenic/non-isogenic pair might be considered more important, we assessed the intensity of the observed phenotypic differences by calculating the percentage of the pairs exhibiting differences by more than 25% and more than 50% (Table 2). We noticed that only 10-25% of the iso/non-iso pairs exhibited differences of >50% for fecundity, midgut mitosis and survival to infection, while no pair exhibited differences of >50% for defecation rate, fecal pH and branching in normoxia or hypoxia (Figure 4A). For the six traits that we had performed z-score analysis in wild-type inbred DGRP strains the range of standard deviation was between 4.25 and 5.5. We then correlated this range for each trait with the percentage of isogenic vs. non-isogenic pairs exhibiting significant differences, as well as, the percentage of those that exhibited more that 25% and 50% differences (Figure 4B-D). We found a significant correlation between the z-score range of each trait and the fraction of isogenic/non-isogenic pairs exhibiting >25% difference (R^2^=0.8053, *p*<0.05; Figure 4C) or >50% difference (R^2^=0.7797, *p*<0.05; Figure 4D). Therefore, the more variable the trait from inbred strain to strain, the more likely *UAS-RNAi* lines sharing half or more of their DNA to exhibit differences of >25%.

**Figure 4:**
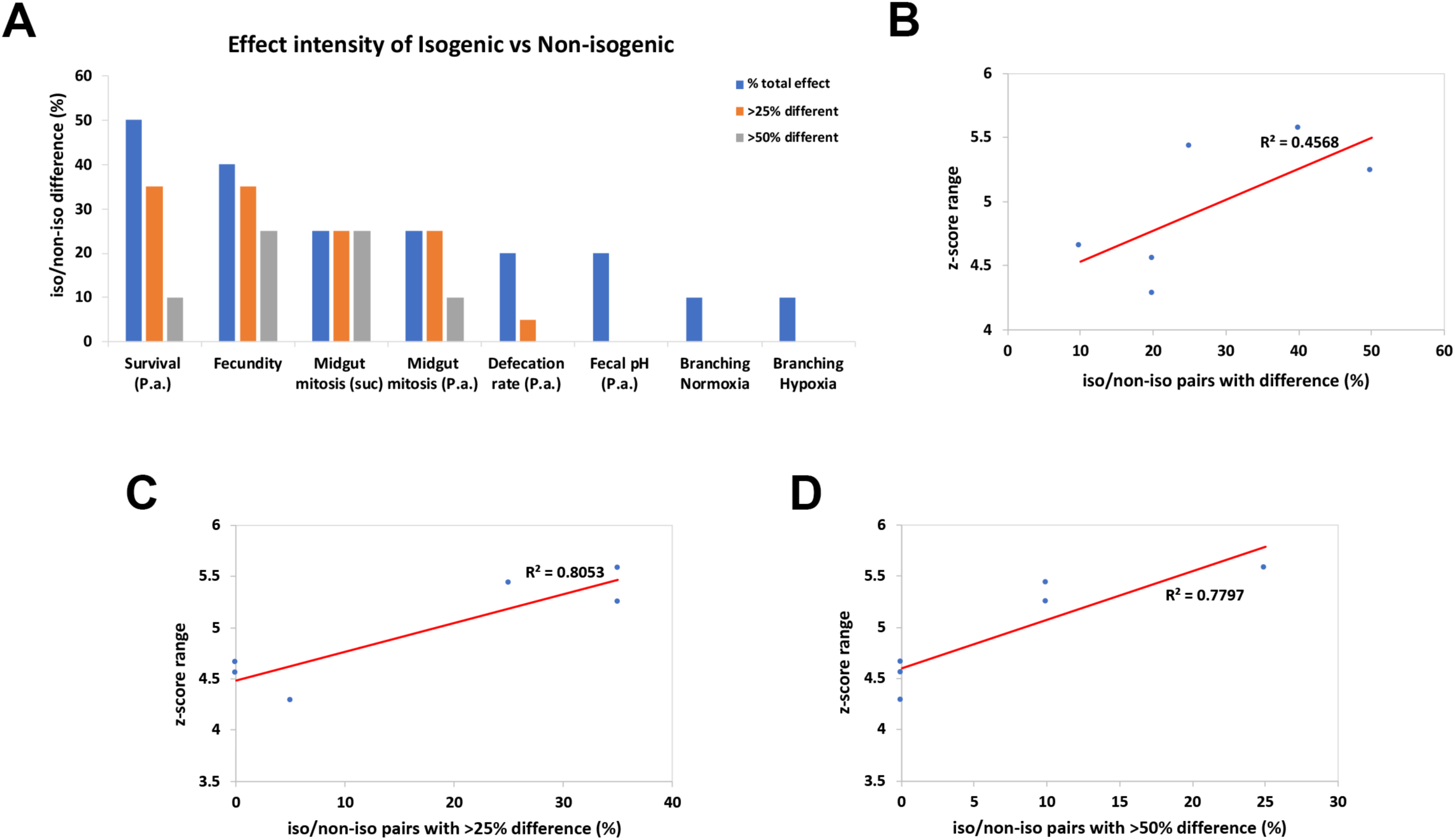
The phenotypic range correlates with the intensity of the difference between isogenic vs non-isogenic phenotypes. (**A**) Percentage of isogenic vs non-isogenic pairs presented with phenotypic differences and subcategories indicating the intensity of the observed differences, more than 25% difference between isogenic/non-isogenic and more than 50% difference between isogenic/non-isogenic. (**B**) Correlation of z-score range of the 6 traits and the percentage of isogenic/non-isogenic pairs that differed significantly for the same traits. (**C**) Correlation of z-score range and the fraction of the total isogenic/non-isogenic pairs that differed by more than 25%. (**D**) Correlation of z-score range and the fraction of the total isogenic/non-isogenic pairs that differed by more than 50%. Trendlines are indicated in red and the R^2^ values are shown in all graphs.

**Table 2:** The intensity of the phenotypic differences in comparisons of isogenic and non-isogenic GD and KK transgenic *UAS-RNAi* lines varies in the absence of Gal4.–

|  | Isogenic vs non-isogenic pairs with difference (20 total) (% of total) | Isogenic vs non-isogenic pairs with >25% difference (% of total) | Isogenic vs non-isogenic pairs with >50% difference (% of total) |
| --- | --- | --- | --- |
| **Fecundity** | 8<br>(40%) | 7<br>(35%) | 5<br>(25%) |
| **Survival upon oral infection** | 10<br>(50%) | 7<br>(35%) | 2<br>(10%) |
| **Midgut mitosis in baseline conditions** | 5<br>(25%) | 5<br>(25%) | 5<br>(25%) |
| **Midgut mitosis upon oral infection** | 5<br>(25%) | 5<br>(25%) | 2<br>(10%) |
| **Defecation rate upon oral infection** | 4<br>(20%) | 1<br>(5%) | 0<br>(0%) |
| **Fecal pH upon oral infection** | 4<br>(20%) | 0<br>(0%) | 0<br>(0%) |
| **Terminal branching in normoxia** | 2<br>(10%) | 0<br>(0%) | 0<br>(0%) |
| **Terminal branching in hypoxia** | 2<br>(20%) | 0<br>(0%) | 0<br>(0%) |

### The genetic background of UAS-RNAi transgenes affects survival upon oral infection in the presence of Gal4

To assess the effect of the genetic background of *UAS-RNAi* lines in the context of Gal4 activation, we crossed the ubiquitously-expressed *act5C-Gal4* driver with the isogenic and non-isogenic *UAS-RNAi* lines. We focused on the trait we found being mostly affected by the genetic background, survival upon oral bacterial infection. We assessed survival of *P. aeruginosa* orally infected flies expressing 11 isogenic vs. their corresponding 11 non-isogenic VDRC *UAS-RNAi* lines (6 GD and 5 KK iso/non-iso pairs) under the control of the ubiquitous *act5C-Gal4* driver. We found significant differences in survival in 4 out of the 11 iso/non-iso pairs crossed to *act5C-Gal4* (Figure 5A). Interestingly, none of the 5 iso/non-iso KK pairs tested exhibited differential phenotypes, when induced by *act5C-Gal4*, although 3 of those exhibited significantly different survival in the absence of Gal4 (Figure 3D). Furthermore, 4 of the 6 iso/non-iso GD pairs tested exhibited significant differences in survival when induced by *act5C-Gal4* (Figure 5B-E). Although 3 of those iso/non-iso GD pairs (45054/GD, 13326/GD, 19172/GD) also affected survival at significantly different levels in the absence of Gal4 (Figure 3C), all 3 behaved the opposite way upon Gal4 induction (e.g. 45054/GD iso exhibited increased and reduced survival compared to 45054/GD non-iso in the presence and absence of *act5C-Gal4*, respectively). Thus, activation of the isogenic vs. the non-isogenic *UAS-RNAi* transgenes via Gal4 leads to differential effects in survival that cannot be predicted by the behavior of the same transgenic lines in uninduced conditions.

**Figure 5:**
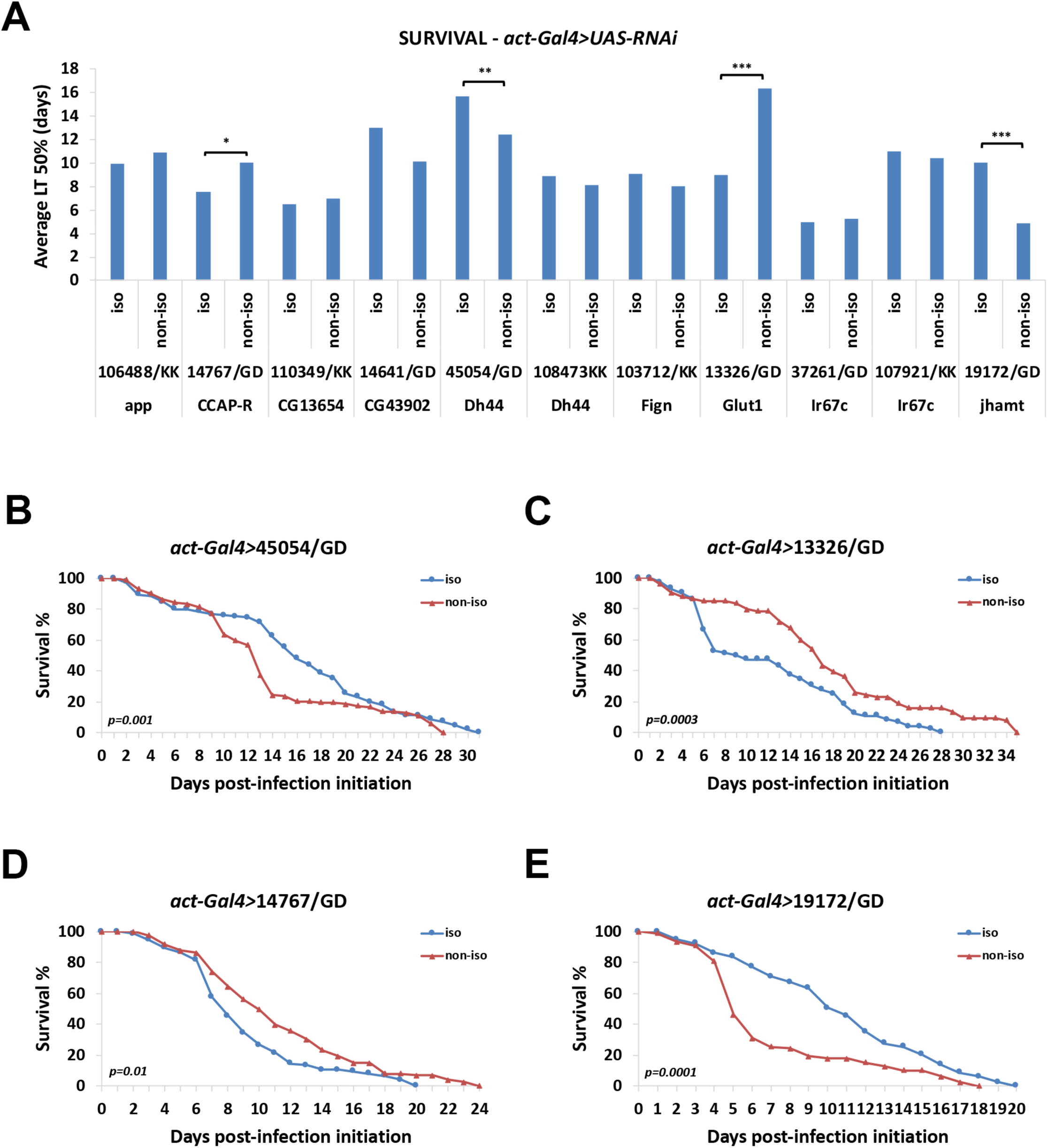
The genetic background affects survival upon Gal4-induction of the transgenic RNAi lines. (**A**) Average LT50% in days upon *P. aeruginosa* infection of *act-Gal4>UAS-RNAi* isogenic vs non-isogenic flies. (**B-E**) Survival curves of the statistically-different isogenic vs non-isogenic *UAS-RNAi* driven by *act-Gal4*. 2-3 independent survival experiments were averaged and *p*-values were calculated by the Kaplan-Meier method (A-E). * 0.01<*p*≤0.05; ** 0.001<*p*≤0.01; *** *p*≤0.001.

### The genetic background affects Gal4 transgenes and midgut dysplasia induced by the same UAS-apc^RNAi^ line

To assess the effect of the genetic background on Gal4 activity per se, we jointly introgressed two transgenes, *UAS-srcGFP* and *Dl-Gal4 (Dl-Gal4>GFP)*, in 22 different DGRP lines. *Dl-Gal4>GFP* allows the visualization of intestinal stem cells in the adult *Drosophila* intestine (Zeng et al., 2010). In addition, it provides a way to assess the generation of dysplastic stem cell clusters (Biteau et al., 2008; Apidianakis et al., 2009) upon genetic perturbations (e.g. by driving expression of UAS lines impinging on mitosis and cell differentiation), during aging or bacterial challenge. After six backcrossing generations to each DGRP strain, the flies were isogenic in the 22 DGRP backgrounds, while heterozygous for the *UAS-srcGFP* and *Dl-Gal4* transgenes. We assessed dysplastic cluster formation (clusters of more than 5 *Dl-Gal4>GFP* positive cells) in these strains by visualizing the GFP in dissected midguts. We found that there was variability in cluster formation in the different homozygous DGRP genotypes (Figure 6A). To assess the effect of overexpression with Gal4, we used an RNAi line targeting *apc* (*UAS-apc^RNAi^*), which, when induced, promotes intestinal stem cell proliferation (Cordero et al., 2012). The phenotypic range was manifold and comparable among: (i) the 22 extensively inbred DGRP lines bearing a single copy of *Dl-Gal4>GFP* (Figure 6A); (ii) the corresponding 22 uninduced *UAS-apc^RNAi^* heterozygotes arising from crosses between a single *UAS-apc^RNAi^* line with each of the 22 DGRP (*Dl-Gal4>GFP*) lines (Figure 6B); and (iii) the corresponding 22 induced and infected *UAS-apc^RNAi^* heterozygotes arising from the same crosses as in (ii) (Figure 6C). Strikingly, no pairwise strain correlation was observed between setups (i) and (ii) (Figure 6D) or between setups (ii) and (iii) (Figure 6E). Thus, the genetic background matters even when the same *UAS-RNAi* line is crossed with Gal4 lines in different genetic backgrounds; and its impact is unpredictable and depends on the specific genetic and experimental setup.

**Figure 6:**
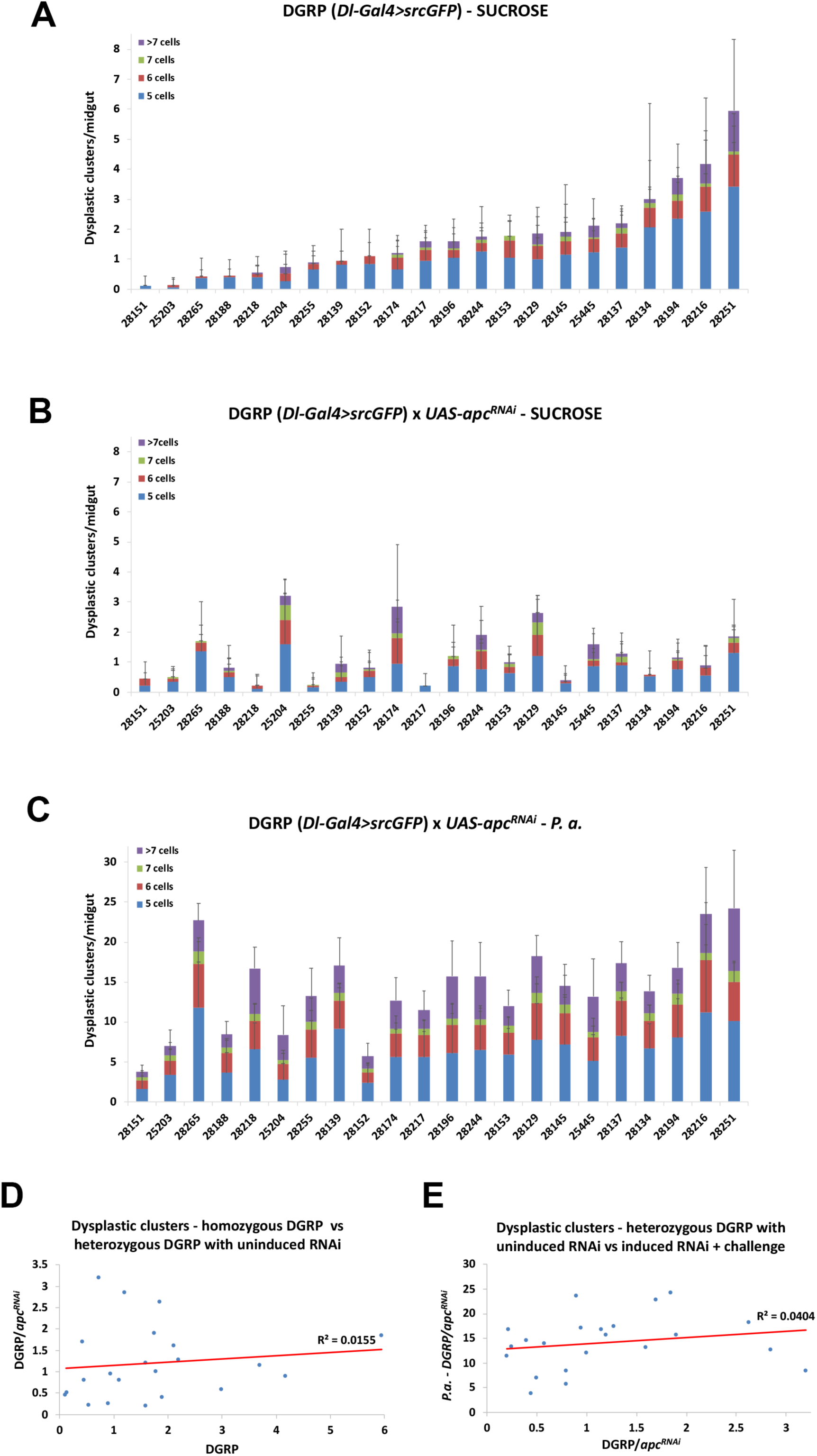
Gal4 is affected by the genetic background. (**A**) Dysplastic cluster formation in 22 DGRP lines carrying one copy of the *Dl-Gal4>GFP* transgenes. (**B**) Dysplastic cluster formation in the progeny of 22 DGRP lines carrying one copy of the *Dl-Gal4>GFP* transgenes crossed to *UAS-apc^RNAi^* in uninduced conditions. (**C**) Dysplastic cluster formation in 22 DGRP lines carrying one copy of the *Dl-Gal4>GFP* transgenes crossed to *UAS-apc^RNAi^* in induced conditions plus oral infection with *P. aeruginosa*. (**D**) Correlation of homozygous DGRP (*Dl-Gal4>GFP*) strains and heterozygous DGRP *Dl-Gal4>GFP/UAS-apc^RNAi^*. (**E**) Correlation of heterozygous DGRP *Dl-Gal4>GFP/UAS-apc^RNAi^* uninduced and induced with oral *P. aeruginosa* infection.

### A modulator of terminal branching lies on the 4^th^ chromosome

A strong genetic background effect from the 4^th^ chromosome was observed during assessment of branching in the isogenic/non-isogenic experiment. Specifically, since terminal branching needed to be assessed in larvae instead of adults, and because the KK VDRC lines had a deep red eye color as heterozygotes, initially, we maintained the isogenized KK strains as heterozygotes and we kept selecting subsequent generations by eye color. Because the use of KK heterozygotes would prevent us from being certain for the genotype of the progeny larvae upon crossing to *w^1118^*, we used a compound balancer stock to ensure that the genetic composition of the progeny would be the desired (Figure 7A). By pre-crossing the heterozygous isogenic KK lines to *yw/Y; T(2;3)/eyFLP2* males, we acquired males with balanced X, 2^nd^ and 3^rd^ chromosomes (but not 4^th^). When these males were crossed to *w^1118^* females, the genotype of the progeny was selected to be isogenic KK for all chromosomes, except for half of the cases of the 4^th^ chromosome (which was derived from the compound balancer stock). Impressively, we noticed that when these KK [via T(2;3)] isogenic strains were crossed to our laboratory *w^1118^*, their progeny differed from those derived from crosses of the non-isogenic strains to our laboratory *w^1118^* in 40% of the cases (4 out of 10 iso/non-iso pairs; 1 down and 3 up in isogenic) in normoxia and in 100% of the cases (10 out of 10 iso/non-iso pairs; all 10 up in isogenic) in hypoxia (Figure 7B-C). Thus, the 4^th^ chromosome of the compound balancer stock imposed a strong enhancing effect on branching.

**Figure 7:**
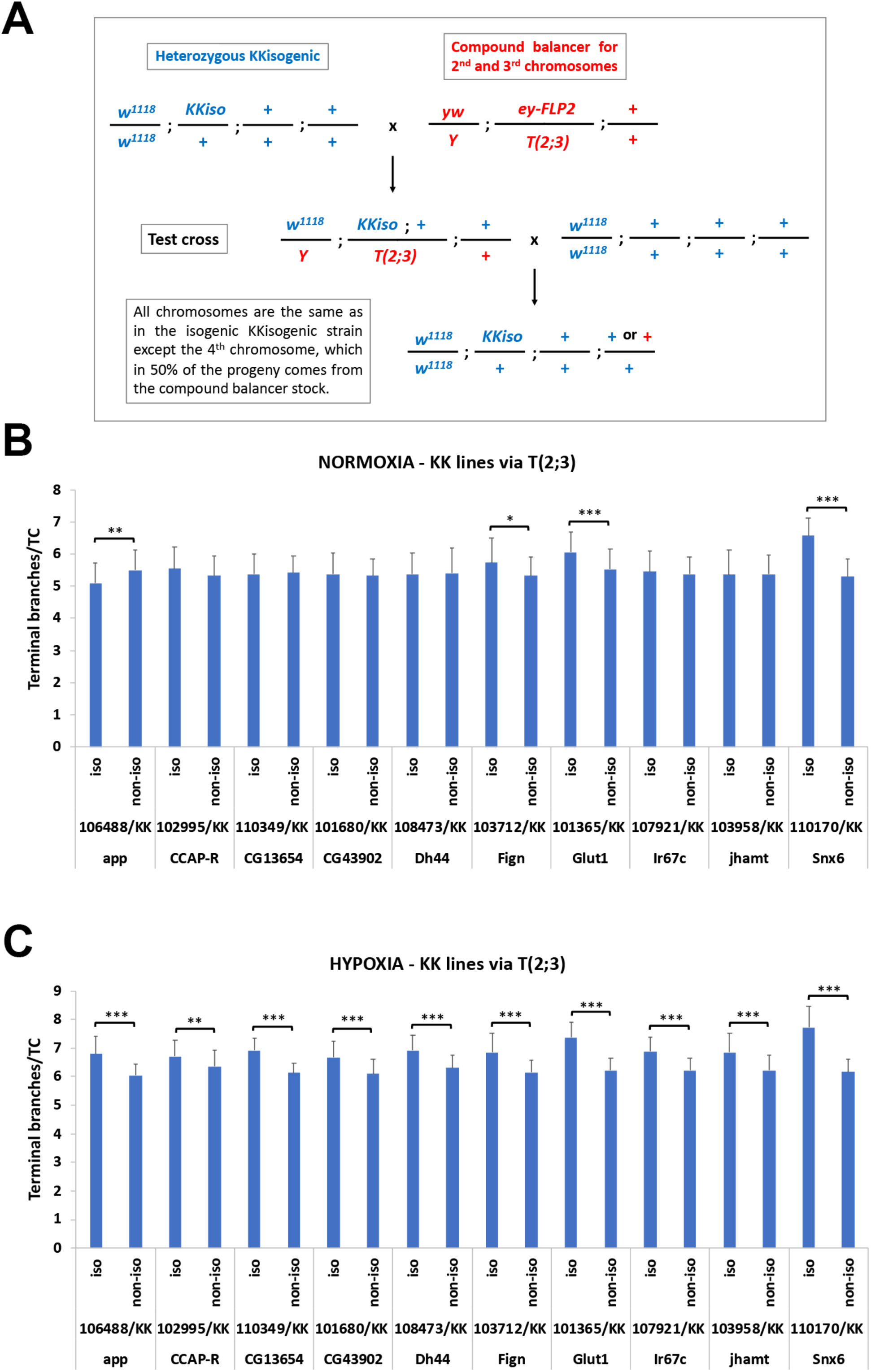
A modulator of terminal tracheal branching lies on the 4^th^ chromosome. (**A**) Crossing scheme used to ensure that larvae assessed for terminal branching carried half of the KK isogenic genotype. Chromosomes X, 2 and 3, but not 4, were always derived from the KK isogenic parent. (**B**) Isogenic via T(2;3) vs non-isogenic comparisons for tracheal branching in normoxia. (**C**) Isogenic via T(2;3) vs non-isogenic comparisons for tracheal branching in hypoxia. Significant pair-wise comparisons are indicated with brackets, whereby * 0.01<*p*≤0.05; ** 0.001<*p*≤0.01; *** *p*≤0.001.

## DISCUSSION

Genetic background effects have been observed in various model organisms ranging from bacteria to mice and in various fitness, physiology and disease phenotypes (Chandler et al., 2013). In this study, the effects of the genetic background on different quantitative traits were primarily measured in inbred homozygous DGRP strains. Not surprisingly, we found that all traits studied (female fecundity, survival upon pathogenic infection, intestinal mitosis upon pathogenic infection, defecation rate and fecal pH upon pathogenic infection and terminal branching in hypoxia) produced variable phenotypes. Based on the genomic information of the DGRP strains, we have identified genomic variants (SNPs, Insertions, Deletions) with a potential role on these phenotypes. Genetic characterization of these variants will be presented elsewhere.

In addition, we observed significant and variable genetic background effects when we assessed transgenic *UAS-RNAi* and *Gal4* constructs in different wild type genetic backgrounds. Specifically, although the VDRC *UAS-RNAi* lines were generated in an isogenic *w^1118^* host strain and, whenever necessary, the balancers used were isogenic to the host strain (Dietzl et al., 2007), when the constructs were introgressed in our laboratory *w^1118^* strain, they produced quantitatively different phenotypes in various traits, when compared to the original, referred to as “non-isogenic”, VDRC lines. This would mean that either the original lines changed over time or that the two *w^1118^* strains (that of VDRC and ours) were not the same, because they were maintained in different laboratory environments. We did not observe major differences between the GD and KK types of transgene insertions in terms of the number of strains affected by isogenization, although the first were generated in random genomic locations using P-element transformation, and thus, cannot be controlled for position effects. This indicates that, not only each GD, but also each KK UAS transgene, which is inserted into the same genomic locus, lies in a variable genetic background. Interestingly, the percentage of isogenic/non-isogenic *UAS-RNAi* pairs exhibiting differences varied for each trait studied. Survival upon oral infection and female fecundity, were more sensitive to genetic background differences with 50% and 40% of the isogenic/non-isogenic pairs exhibiting significantly different phenotypes. Only 10-25% of the isogenic/non-isogenic pairs exhibited differences of >50% for fecundity, midgut mitosis and survival to infection, whereas no pair exhibited differences of >50% for defecation rate, fecal pH and branching in normoxia or hypoxia. Moreover, the extent to which each trait varied among the ~150 inbred strains correlated with the percentage of isogenic/non-isogenic pairs differing for the same trait, but only when the differences when >25%. Thus, the more variable the trait from strain to strain, the more likely *UAS-RNAi* lines sharing half or more of their DNA to exhibit differences of >25%.

Gal4-induction of the isogenic and non-isogenic *UAS-RNAi* transgenic strains was used to assess the trait affected the most by the genetic background, survival upon oral infection. Although we were able to assess only 11 isogenic/non-isogenic pairs upon *act5C-Gal4* induction, we found that 4 out of those exhibited significant difference in survival. Interestingly, all pairs presented with differential survival were GD strains. Nevertheless, none of the 3 isogenic/non-isogenic pairs that exhibited differential survival upon infection could be predicted based on its phenotype in uninduced conditions (Figure 5 and Figure 3C). Furthermore, Gal4-dependent phenotypes were also observed, in inbred strains, as well as upon crossing of those to the same *UAS-RNAi* strain and environmental challenge (Figure 6). Similarly, the observed differences upon induction could not be attributed to the initial differences of the inbred strains. Thus, Gal4/UAS genotypes sharing half of their DNA exhibit differences that cannot be predicted based on prior knowledge on a similar genetic setup.

Moreover, we found a strong effect from the 4^th^ chromosome in terminal branching, when assessing isogenic genotypes through balancing via a compound balancer strain and a genetic scheme ensuring that X, 2 and 3 chromosomes derived from the KK isogenic, but the 4^th^ chromosome was 50% that of the KK isogenic and 50% that of the balancer stock (Figure 7A). Specifically, the compound balancer 4^th^ chromosome increased branching. Since the 4^th^ chromosome is small, largely heterochromatic and encompasses approximately 100 genes (Riddle and Elgin, 2018), it would be interesting to assess if any of these plays a role in terminal branching. Initial Gene Ontology (GO) analysis of the 4^th^ chromosome genes, indicates that three of them (*ci*, an effector of the Hh pathway, *pan*, an effector of the Wg pathway, and *zyx*, a cytoskeletal protein) function in the trachea and *zyx* is involved in terminal branching morphology and tracheal air filling (Merabet et al., 2005; Renfranz et al., 2010).

Since the genetic toolkit of *Drosophila* is continuously expanding, we tend to use more and more transgenic lines to understand the function of genes of interest. This is especially true for *UAS-RNAi*, that allows tissue-specific knockdown, when combined with *Gal4* (Mohr and Perrimon, 2012; Kaya-Çopur and Schnorrer, 2016). Depending on the phenotype we aim to study, we should be aware that the genetic background matters more or less. For example, highly variable (multi-organ or multi-factorial) traits, such as survival and fecundity, are seriously affected by the genetic background, whereas less variable traits, such as tracheal branching, are much less affected. So, what should one do to ensure that the phenotypes produced in his/her screen are not because of background effects? Considering the findings of this study, we propose the following guidelines:

1. Isogenization of all transgenic lines used in a study in the same genetic background is ideal and allows direct comparisons of strains, but it is time-consuming. Some researchers recommend using more than one wild-type background for isogenization. This is feasible and desirable, when one’s study focuses on a particular gene.
2. If isogenization is impractical, as in cases of large-scale screening of *UAS-RNAi* lines, only differences >50% for highly-variable traits or >25% for less variable traits can be considered. This is because only 10-25% of the isogenic/non-isogenic pairs exhibited differences of >50% for highly-variable traits and because isogenic/non-isogenic pair differences of >25% correlated with the variation of the phenotypic (z-score) range of wild-type flies. Accordingly, one should know one’s assay and the range of phenotypic differences expected. A small-scale pilot experiment can provide essential information of the phenotypic range. For example, terminal tracheal branching has a narrow phenotypic range and, thus, smaller phenotypic differences might be significant.
3. Experiments must be repeated independently three times to account for the impact of gene-to-environment variation.
4. Validation of an observed phenotype must be performed in multiple ways, e.g. when assessing RNAi phenotypes, multiple genetically-different *UAS-RNAi* lines targeting the same gene must be used. These might be obtained from different sources (stock centers or independent laboratories) and ideally should target independent regions of the gene of interest.

## MATERIALS AND METHODS

### Drosophila maintenance

All strains and crosses were maintained on standard agar cornmeal fly food in a 12-hour light-dark cycle in a temperature-controlled incubator (Fitotron) at 25°C (unless specified otherwise) with 65% humidity. For the DGRP screening for fecal spot number and pH measurements, prior to infection, female mated flies were aged for 4 days at 25°C in bottles changed daily containing fly food supplemented with 50 ug/ml of the broad-range antibiotic Rifampicin, which does not kill PA14, but can eliminate most of the microorganisms present in the intestine of the flies. Rifampicin was also used in screening of the isogenic/non-isogenic pairs (Figure 3) for all assays except branching.

### Drosophila Strains

The *Drosophila* Genetics Reference Panel (DGRP) collection of wild type inbred sequenced strains (Mackay et al., 2012; Huang et al., 2014) was used in all screens assessing physiological traits (fecundity, survival, intestinal regeneration, fecal motility, fecal spots, tracheal branching). The Vienna *Drosophila* Research Center (VDRC) UAS-RNAi lines used in this study are the following: 32863/GD and 106488/KK targeting *CG42318/app*, 45054/GD and 108473/KK targeting *CG8348/Dh44*, 19172/GD and 103958/KK targeting *CG17330/jhamt*, 13550/GD and 110349/KK targeting *CG13654*, 14641/GD and 101680/KK targeting *CG43902*, 14747/GD and 102995/KK targeting *CG33344/CCAP-R*, 37261/GD and 107921/KK targeting *CG32058/Ir67c*, 24275/GD and 110170/KK targeting *CG8282/Snx6*, 24746/GD and 103712/KK targeting *CG3326/Fign*, 13326/GD and 101365/KK targeting *CG43946/Glut1*. Other stocks used in this study were the following (source and/or stock center numbers in parentheses): *UAS-apc^RNAi^* (VDRC, #1333), *UAS-srcGFP* (BDSC, #5432), *Dl-Gal4* (Zeng et al., 2010), *yw; ey-FLP2/T(2;3)CyO;TM6B,Tb,Hu* (BDSC, #8204), *actin5C-Gal4* (BDSC, #25374).

### Isogenization of transgenic lines

The VDRC lines, referred to as “non-isogenic”, were backcrossed to the laboratory *w^1118^* strain for at least 6 generations to produce the “isogenic” lines. For the GD lines, the transgenes were homozygosed following the backcrossing based on the intensity of their eye color. For the KK lines, due to the intensity of the red eye color, it was safer to maintain the transgenes in a heterozygous state (unless specified otherwise). Since a fraction of the KK strains has been shown to carry two inserts (Green et al., 2014; Vissers et al., 2016), we tested all of them during the isogenization process for segregation of different eye colors, as well as by PCR (Green et al., 2014); we found that 101680/KK and 103958/KK carried double insertions (data not shown). Nevertheless, we did not observe any bias in iso/non-iso variation between these particular pairs.

### Fecundity measurement assays

Female flies were allowed to mate for 2 days at 25°C. Subsequently, males are removed from the cultures and females are left to lay eggs for 24 hours in a square bottle with 50 ml of food. Females are flipped to a new bottle for egg-laying every day for the next 3 days. Average fecundity is calculated as the number of progeny produced by the females in the 4 bottles divided by the number of females.

### Pseudomonas aeruginosa infection

A 3 ml overnight culture of *Pseudomonas aeruginosa* PA14 was diluted 1:100 in LB (Luria Bertani medium) to prepare an overday 3 ml culture. When the culture reached OD_600_=3, it was used to prepare the infection mix (5 ml per vial: 3.5 ml ddH_2_O, 1 ml 20% sucrose and 0.5 ml PA14 OD_600_=3) or sucrose mix (5 ml per vial control: 4 ml ddH_2_O and 1 ml 20% sucrose). 5 ml of the infection or control mix was used to soak a cotton ball in a narrow fly vial, which was subsequently plugged with a dry cotton ball. After 4-5 hours starvation in empty vials, the flies were put in the infection vials and incubated at 25°C with the cotton plug facing down.

For the fecal spot and pH measurements, flies were fed concentrated bacteria of OD_600_=50 (when the overday culture reached OD_600_=2, it was concentrated 25 times by centrifugation and resuspension of the bacterial pellet in 4% sucrose). Specifically, each feeding vial plugged with a cotton ball contained 5 ml agar gel (3% w/v agar in H_2_O) on top of which a Whatmann disc with 200 ul of the infection mix was placed. 25 mated starved young female flies were allowed to feed for 15 hours at 25°C on infection mix (or 4% sucrose for control) in each vial and subsequently their fecal spots were assessed for numbers and pH.

### Survival assays

Young female flies were subjected to intestinal infection with the virulent *Pseudomonas aeruginosa* strain PA14. The percentage of dead flies was calculated daily as the (number of dead flies per vial/total number of flies) × 100 until all flies are dead in each vial. LT50% (lethal time 50%), the time when 50% of the flies were dead was used as an indicator of survival for comparisons. For *act5C-Gal4* experiments, the crosses with the UAS-RNAi strains were maintained at 18°C to suppress expression of the RNAi. Emerging female flies were allowed to mature and mate at 25°C and were subsequently subjected to infection with PA14 at 25°C, as described above.

### Intestinal regeneration measurement

Female adult flies (4-7 days old) were subjected to oral infection with the pathogenic strain of *Pseudomonas aeruginosa* PA14. Five days upon infection initiation, intestinal mitosis was measured by staining the intestines with the Rabbit-anti-pH3 antibody (Millipore) followed by secondary detection with Donkey-anti-Rabbit Alexa555 (Invitrogen), as described previously (Apidianakis et al., 2009). Mitotic cells were counted in whole midguts under the fluorescent Zeiss Axioscope A1. At least 10 midguts per genotype were used for mitotic index calculations per genotype.

### Fecal number and pH measurements

Infected female flies were starved in empty vials for 5 hours and then placed in vials with cotton balls impregnated with 5 ml 4% sucrose with 0.5% w/v bromophenol blue (BPB) pH=7. BPB is a pH-sensitive dye, which is yellow in pH<3 and turns blue in pH>4.7. 50 flies were split in 3 vials with BPB and were allowed to feed for 5 hours (vials were placed in the incubator with the cotton plug facing down). 10 flies from each vial were moved to a petri dish containing a sterile cotton ball impregnated with 2.5 ml of the BPB solution and were left to poo for 20 hours. Then, the flies were removed from the plates and the total number of fecal spots was measured in the 3 plates. After counting the fecal spots, 10 spots from 2 separate locations of each plate (6 samples) were collected in an microtube in 25 ul ddH_2_O pH=5.5 and their pH was measured with a fine tip pH meter. All experiments were performed at 25°C.

### Terminal tracheal branching

Vials containing larvae were capped with net during development to allow quick gas exchange and were subjected to hypoxia (5% O_2_) for 4 hours in a controlled chamber in the 25°C incubator. For control experiments, larvae were maintained in normoxia (21% O_2_) in the 25°C incubator. Late L3 larvae were aligned on a microscope slide (dorsal side up) and heat-killed by placing the slide on a 70°C hot plate for 3-5 sec (Ghabrial and Krasnow, 2006). Subsequently, and in less than one hour upon heat-killing, the two terminal tracheal cells of the second tracheal metamere were assessed for the number of their terminal branches and photographed under bright-field on a Zeiss Axioscope A1.

### Dysplastic cluster assays

Dysplastic clusters encompassing more than 5 cells were counted in adult *Drosophila* midguts of flies expressing GFP in intestinal stem cells via *Dl-Gal4>GFP*. *UAS-srcGFP* and *Dl-Gal4* were introgressed together in 22 DGRP lines. After six backcrossing generations, the transgenes were maintained in the DGRP backgrounds by selection of GFP expression and were kept in a heterozygous state. Baseline cluster formation was measured in 5-7 days old adult females maintained at 25°C. For *apc^RNAi^* experiments, the DGRP (*Dl-Gal4>GFP*) isogenic strains were crossed to *UAS-apc^RNAi^* at 25°C and the adult female progeny were aged for 3 days at 25°C before dysplastic cluster measurement for the uninduced state (the fact that no increase of cluster number was observed confirmed that *apc^RNAi^* was not induced in these conditions). For *apc^RNAi^* induction, the adult female progeny were aged for 3 days at 25°C before feeding with *P. aeruginosa* PA14 for 2 days at 29°C, where *apc^RNAi^* is induced concomitantly with infection, which boosts the regenerative ability of the gut. Dysplastic clusters were measured under a fluorescent Axioscope A.1 microscope (Zeiss) in dissected midguts (Apidianakis et al, 2009) upon fixation and staining with rabbit- or chicken-anti-GFP (Invitrogen) and mouse-anti-Prospero (DSHB) followed by secondary detection with anti-rabbit or anti-chicken Alexa 488 and anti-mouse Alexa 555 (Invitrogen).

### Statistical analysis

Normalized z-scores for each DGRP screen were calculated by the formula z=(x−μ)/σ, where x is the mean observed value for each genotype normalized to the set average (each DGRP screen was performed in multiple sets of at least 10-20 genotypes), μ is the normalized population mean and σ is the standard deviation of the population. For hypoxia branching, z-score was calculated from the observed genotype means without normalization of the values, because the genotypes assessed per set were few.

For the isogenic/non-isogenic pair screening, initially each experiment was performed in duplicate and if both times the result had the same trend (at least once significant based on Student’s t-test), the experiment was repeated for a third time. If 2 out of 3 times the result was statistically significant with the same trend, the difference was considered significant (noted with p-values in Figures 3 and 7, * 0.01<*p*≤0.05; ** 0.001<*p*≤0.01; *** *p*≤0.001).

For survival assays the p-value was calculated with the Kaplan Meier estimator using the log-rank test (MedCalc statistical software). For all other assays, p-values were calculated with the Student’s t-Test in Excel (Office package). For correlation graphs, trendlines and R^2^ values were calculated in Excel and *p*-values were calculated by Pearson *r* value and the number of values correlated.

## ACKNOWLEDGMENTS

The authors would like to thank Maria Glykainou, Patapios Patapiou and Styliani Irakleous for help with DGRP screening and Androniki Giakoumi for help with *act-Gal4* survival assays. We also thank the VDRC and BDSC for the *UAS-RNAi* and DGRP fly strains used in this study, respectively, the DSHB for antibodies, and Steven Hou for the *Dl-Gal4* stock. Work in the Pitsouli and Apidianakis laboratories was supported by the Marie Curie GIG FP7 - People and the Fondation Santé.

